# A general strategy to engineer high-performance mammalian Whole-Cell Biosensors

**DOI:** 10.1101/2024.02.28.582526

**Authors:** Alessio Mallozzi, Virginia Fusco, Francesco Ragazzini, Arne Praznik, Roman Jerala, Diego di Bernardo

## Abstract

Transcription-based whole-cell biosensors (WCBs) are cells engineered with an analyte-responsive promoter driving the transcription of a reporter gene. WCBs can sense and report on bioactive molecules (analytes) relevant to human health. Designing an analyte-sensitive promoter requires a cumbersome trial-and-error approach and usually results in biosensors with poor performance. Here, we integrated Synthetic Biology with Control Engineering to design, computationally model, and experimentally implement high-performance biosensors in mammalian cells. Our approach, unlike traditional methods, does not rely on optimizing individual components such as promoters and transcription factors. Instead, it uses biomolecular circuits to enhance the biosensor’s performance despite inherent component flaws. We experimentally implemented eight different biosensors by employing CRISPR-Cas systems, then quantitatively compared their performance and identified one configuration, which we named CASense, that overcomes the limitations of current biosensors. Our approach is generalisable and can be adapted to sense any analyte of interest for which there is an analyte-sensitive promoter, making it a versatile tool for multiple applications. As a proof of concept, we engineered a high-performance biosensor of intracellular copper due to the critical role that copper plays in human health and disease, and to the lack of techniques able to measure intracellular copper levels in living cells. The significance of our work lies in its potential to impact the monitoring of bioactive molecules and chemicals both in vitro and in vivo, which is crucial in areas such as toxicology, drug discovery, disease diagnosis and therapy.

## Introduction

Mammalian whole-cell biosensors (WCBs) are cells engineered with biomolecular circuits providing them with the ability to sense and report on bioactive molecules, chemicals, or even pathogens (analytes) relevant to human health that are present within the cell or in the surrounding microenvironment [1, 2, 3, 4]. Beyond sensing, WCBs can be coupled to effector proteins to control a biological process of interest. Applications range from toxicology and drug discovery, to biomanufacturing and biosecurity. WCBs are thus attractive tools in biotechnological and biomedical applications. However, only a handful of WCBs are currently available, since developing a biosensor requires a cumbersome trial-and-error approach as initial prototypes usually perform poorly [1, 5, 6, 7, 8, 9, 10, 11, 12, 13]. Hence, a general strategy for the design and experimental implementation of biosensors is of paramount importance.

Genetically encoded biosensors can be developed using different architectures according to the way the analyte is sensed. Transcription-based biosensors are illustrated in Figure 1; they consist of an analyte-responsive promoter regulated by an endogenous, or synthetic, transcription factor. The activity of the transcription factor is modulated by the target analyte and results in the reporter gene’s transcription.

**Figure 1:**
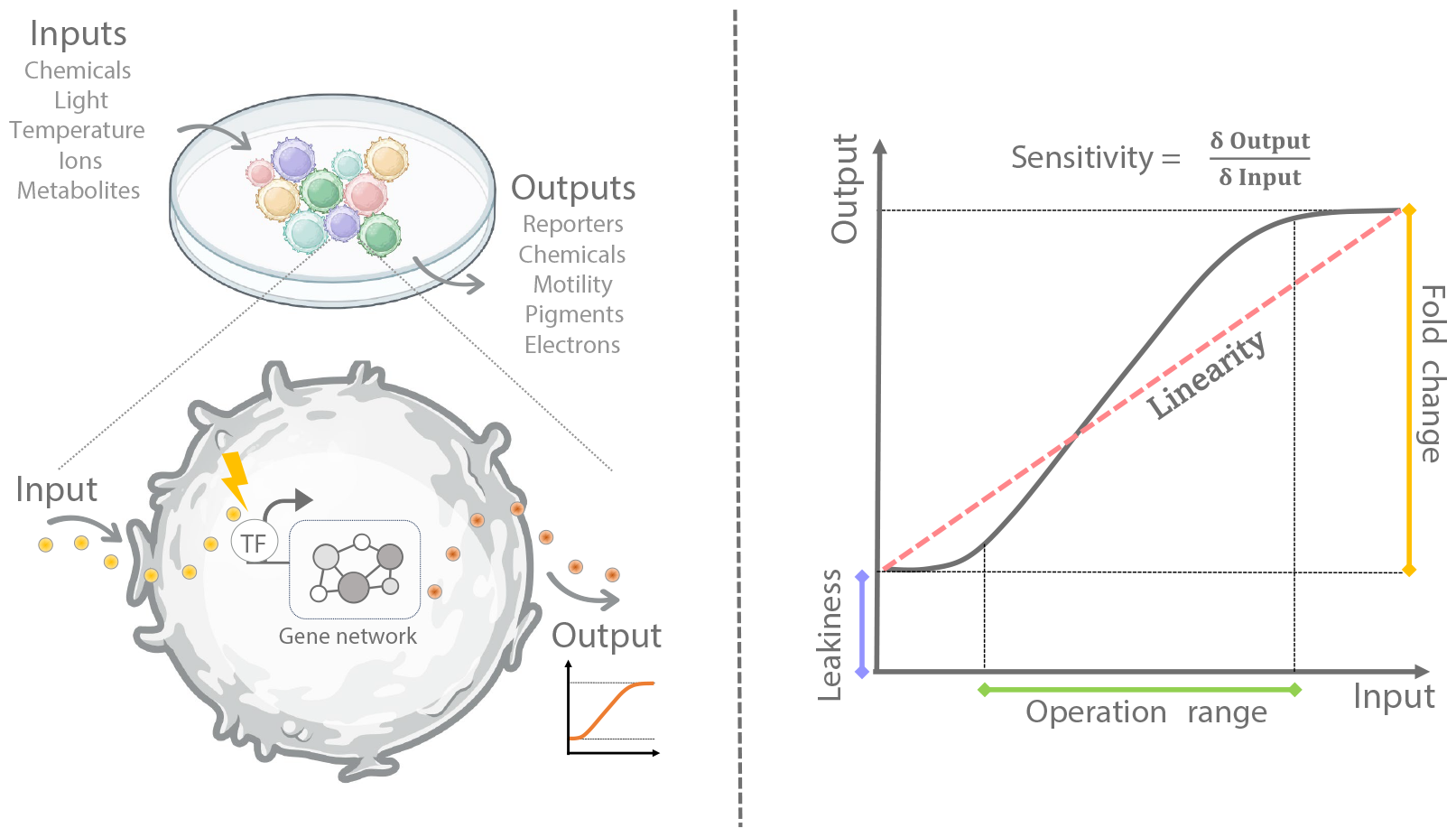
Whole-cell biosensor and its main feautures. Whole-cell biosensors are living cells that can detect and measure the concentration of specific analytes (input) such as heavy metals, by producing a measurable signal (output), such as fluorescence or luminescence. A biosensor can be characterised by five main features: **Leakiness:** the basal expression of the reporter (output) in the absence of the analyte (input); **Fold-Change**: the ratio between the maximum reporter expression in the presence of the analyte and the leakiness; **Sensitivity:** the steepness of the dose-response curve (i.e. its derivative); **Operation Range:** the difference between the maximal and minimal concentration of the analyte that can be detected; **Linearity:** the extent of deviation from a linear dose-response curve, i.e. when the output value is directly proportional to the analyte concentration.

WCBs can be characterised by features to gauge their performance [14], as illustrated in Figure 1. State-of-the-art transcription-based WCBs are not efficient as they exhibit unintended basal expression (“leakiness”) in the absence of the analyte and limited sensitivity, linearity, operation range, and fold change.

In this work, we employed a synergistic approach integrating Synthetic Biology with Control Engineering to yield high-performance transcription-based biosensors in mammalian cells. Our approach, unlike traditional methods, does not rely on optimizing individual components like promoters and transcription factors. It uses, instead, gene networks to enhance the biosensor’s performance by effectively addressing and compensating for inherent flaws in individual components. As proof of principle, we applied our strategy to engineer high-performance biosensors of intracellular copper, by leveraging a previously published copper-responsive promoter [15].

## Results

### Computational investigation of gene network motifs to design high-performance biosensors

The design of high-performance biosensors aims at improving the five features of the input-output response schematised in Figure 1. We thus selected gene network motifs that could yield optimal features [10, 16, 17, 18, 19] and then combined them into more complex gene networks to build high performance biosensors, as depicted in Figure 2. The performance of each gene network was assessed by comprehensive mathematical and computational analyses. We considered at first five biosensors’ designs:

**Figure 2:**
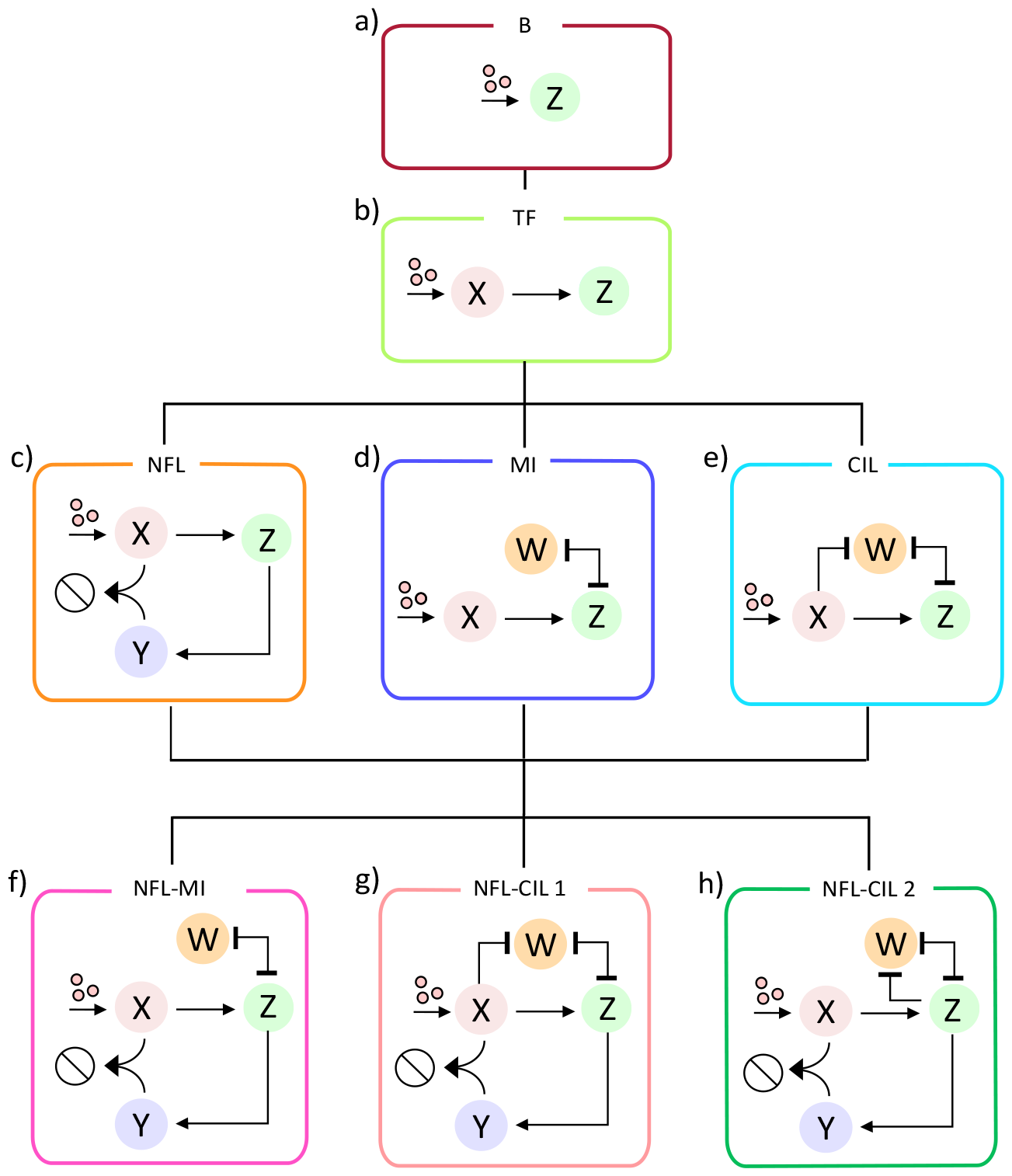
Alternative gene networks to improve the design of a biosensor. **a**, Basic biosensor (B): an analyte–responsive promoter drives the production of the output species *Z*. **b**, Transcription Factor amplified biosensor (TF): an analyte–responsive promoter drives the expression of transcription factor *X* that in turn activates the expression of the output *Z*. **c**, Negative Feedback Loop biosensor (NFL): the TF biosensor is augmented with a repressor species *Y* that sequesters *X* and inhibits its function; the output *Z* activates the expression of *Y*. **d**, Mutual Inhibition biosensor (MI): the TF biosensor is augmented with a constitutively expressed species *W* that can repress, and be repressed by, output *Z*. **e**, Coherent Inhibitory Loop Biosensor (CIL): the MI biosensor is modified so that species *W* can be repressed by species *X*. **f**, NFL-MI biosensor: the NFL biosensor is augmented with the constitutively expressed species *W* of the MI biosensor. **g**, NFL-CIL biosensor version 1: the NFL-MI biosensor is modified so that species *W* can now be repressed by species *X*. **h**, NFL-CIL biosensor version 2: a modification of the NFL-MI biosensor where species *W* can be transcriptionally repressed by species *Z. Pointed arrows represent activation, blunt head arrows represent repression; mutual repression is represented by an arrow with a blunt head and a blunt tail*.

- **Basic biosensor (B)**. This biosensor, shown in Figure 2.a, comprises the gene encoding for the reporter protein, *Z*, placed downstream of an analyte-responsive promoter that responds to changes in analyte concentration.
- **Transcription Factor amplified biosensor (TF)**. The simplest modification to the basic biosensor is to add an exogenous transcription factor (*X*) between the analyte-responsive promoter and the reporter gene *Z* to amplify the output [16, 20, 21], as shown in Figure 2.b.
- **Negative feedback loop biosensor (NFL)**. One of the tenets of Control Engineering is that adding negative feedback can increase the linearity of the input-output response of a dynamical system [10, 18, 19], in addition to increasing its robustness to external disturbances [22]. We thus modified the TF biosensor by adding a species *Y* implementing a negative feedback, as shown in Figure 2.c. In this implementation, protein *Z* controls the expression of the inhibitor *Y* that stoichiometrically binds to the activator *X* thus leading to its inactivation and closing the loop[23].
- **Mutual Inhibition (MI) and Coherent Inhibitory Loop (CIL) biosensors**. These two gene network motifs (Figure 2.d,e) are able to reduce leaky gene expression of the reporter *Z* and to improve sensitivity and fold change [16, 17]. In both networks, the analyte-dependent promoter drives the transcription-factor *X* that activates species *Z*, while species *W* and *Z* mutually repress each other at the post-transcriptional level. The mutual inhibition (MI) motif in Figure 2.d exhibits a constitutively expressed species *W*, while in the Coherent Inhibitory Loop (CIL) in Figure 2.e, species *W* is transcriptionally repressed by species *X*. In both networks, in the absence of the analyte, *X* is not expressed, while species *W* represses *Z*, thus minimising leakiness. On the contrary, in the presence of the analyte, *X* is expressed and induces expression of *Z* leading to repression of *W*. In the CIL motif, *X* also directly represses *W* thus further contributing to maximal expression of *Z*.

We derived and analysed quantitative models based on Ordinary Differential Equations (ODEs) for each of the five gene networks to assess their performance in terms of leakiness, fold change, sensitivity to input, linearity, and operation range, as detailed in Supplementary Material. Computational results are summarised in Supplementary Figure S1. The TF biosensor (Supplementary Figure S1.a) has a worse overall perfomance than the one of the basic biosensor, since it just amplifies the output *Z* by shifting the whole dose-response curve up (Supplementary Material). The NFL biosensor (Supplementary Figure S1.b) improves two out of the five features of the basic biosensor, i.e. the linearity and operation range, as it linearises the dose-response curve. Finally, both MI and CIL biosensors (Supplementary Figure S1.c,d) improve leakiness, fold-change and sensitivity, at the cost of a worsening in linearity and operation range. All together, the computational analysis demonstrates that the NFL provides improvements that are complementary to those conferred by the MI and CIL motifs. We thus reasoned that by combining the NFL with the MI and CIL network motifs, we could improve all of the five features at the same time. To test this hypothesis, we designed, modeled, and analyzed three additional gene networks, as show in Figure 2.f-h:

- **NFL-MI biosensor** The NFL-MI combines the NFL and MI motifs by adding to the NFL motif, the constitutively expressed species *W* that both represses, and is repressed by, species *Z* (Figure 2.f).
- **NFL-CIL version 1** The NFL-CIL version 1 combines the NFL and CIL motifs by adding species *W* to the NFL motif under the control of a promoter that can be transcriptionally repressed by species *X* (Figure 2.g),
- **NFL-CIL version 2** The NFL-CIL version 2 again combines the NFL and CIL motifs, but this time species *W* is driven by a promoter that can be transcriptionally repressed by species *Z* itself (Figure 2.h).

The quantitative analysis of the newly proposed biosensors is reported in Supplementary Figure S1.e-g and reveals that the three networks do improve all of the five features when compared against the basic biosensor, with the NFL-CIL version 2 exhibiting the best performance.

Taken together, our results suggest that it may be possible to implement biosensors with improved performances without requiring to modify the biosensor’s core constituent, i.e. the analyte-sensitive promoter.

### Experimental implementation of the gene networks as whole-cell biosensors

We thus set to experimentally implement the eight gene networks in order to compare their performances and select a biomolecular circuit implementing a high-performance whole-cell biosensor.

The basic copper biosensor in Figure 3.a consists of a firefly luciferase (FLuc) placed downstream of a synthetic promoter (pMRE) containing four metal-responsive elements (MREs)[15]. These MREs serve as binding sites for the endogenous metal-responsive transcription factor 1 (MTF-1). Upon exposure to copper dichloride, zinc ions are displaced from intracellular metallothioneins and bind to MTF-1, leading to its activation and triggering transcription from the pMRE promoter[15]. This promoter has been shown to be specific for copper and zinc, but not to other metals[15]. We transfected a human cell line (Hek293T), which endogenously express MTF-1, with a plasmid encoding the basic biosensor (Materials and Methods) and assessed its performance at increasing doses of copper dichloride, as reported in Figure 3.b. From the dose-response curve, we derived the values of the biosensor’s features, as shown in Figure 3.c, which we used as a baseline for examining the performance of the other gene networks.

**Figure 3:**
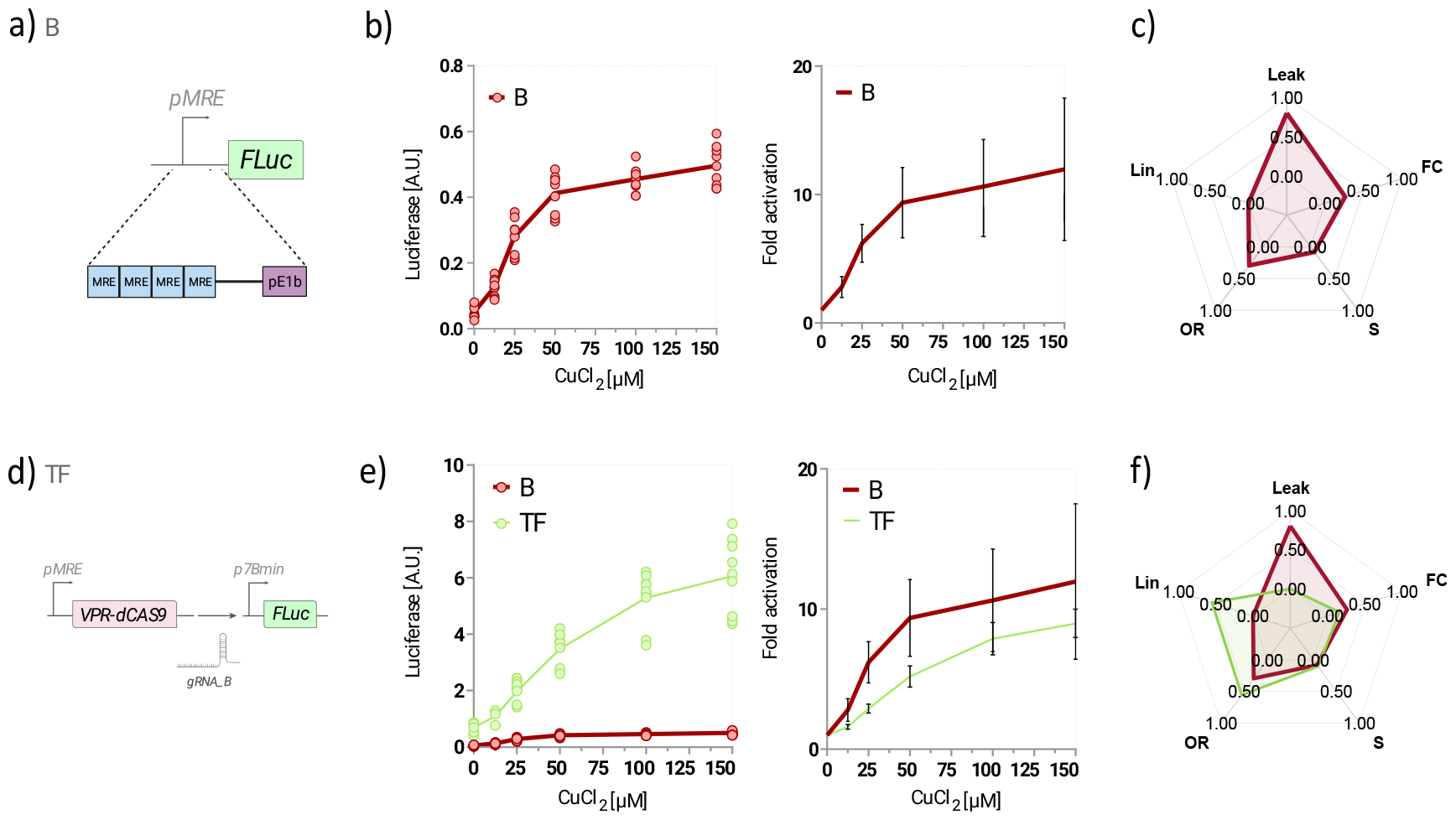
Experimental implementation of the Basic and TF-amplified biosensors. **a,d**, Schematic representation of the constructs impementing the basic and TF-amplified biosensors. **b,e**, Dose-response curve of the Basic biosensor (red) and the TF-amplified biosensor (green) for increasing copper dichloride concentrations. n=9 biological replicates from three independent experiments. Firefly luciferase (FLuc) expression was evaluated by luminescence measurements normalised to Renilla firefly (RLuc) and reported either as Arbitrary Units [A.U.] (left panel), or as Fold Activation obtained by dividing each data point by the average luminescence in the absence of copper (right panel). **c,f**, Radar plot reporting the values of the five features of the Basic biosensor (red) and of the TF-amplified biosensor (green). Leak: leakiness, FC: fold change, S: sensitivity to input, OR: operation range, Lin: linearity. The score ranges from 0, which is assigned to the biosensor with the worst performance, to 1 for the one with the best performance. ***VPR-dCas9***: *nuclease-deficient Cas9 fused to the transactivation domain VPR;* ***FLuc***: *firefly Luciferase;* ***gRNA***_*B: gude RNA with sequence B;* ***pMRE***: *synthetic promoter harbouring four metal responsive elements;* ***p7B_min***: *a synthetic promoter consisting of 7 sequences complementary to gRNA_B upstream of a minimal CMV promoter*.

As a further control of intracellular copper concentration, we used Copper Fluor-4 (CF4), a recently developed a Cu^+^-specific fluorescent sensor, which is stable in a physiologically relevant pH regime and suitable for live-cell imaging[24]. We quantified CF4 fluorescence at increasing concentration of copper dichloride. The resulting dose-response curve in wild-type HEK293T cells is reported in Supplementary Figure (Supplementary Figure S2) and has a behaviour similar to the one of the basic biosensor.

To implement the TF-amplified biosensor shown in Figure 3.d, we chose as species *X*, the nuclease-deficient Cas9 fused to the transactivation domains VP64, p65, and Rta (VPR) (VPR-dCas9) [25] driven by the synthetic copper-responsive promoter pMRE; species *Z* is the FLuc driven by the synthetic promoter, p7B Min, which is strongly activated by the VPR-dCas9 in the presence of the guideRNA gRNA_B [26]. The dose-response curve for increasing concentration of copper dichloride is depicted in Figure 3.e and demonstrates that the TF biosensor amplifies luciferase expression with respect to the basic biosensor, as expected. This amplification however results in a worsening of the leakiness, in agreement with the computational results, while leaving the other features unchanged except for a minor improvement in linearity, as reported in Figure 3.f

### Experimental implementation of the NFL motif

The experimental implementation of the NFL motif required to modify the TF biosensor in Figure 3.d with an additional species *Y* able to inactivate the synthetic transcription factor VPR-dCas9. We thus chose as species *Y* the Anti-CRISPR protein AcrIIA4, which is able to bind to and inactivate the cognate Cas9 [27, 28].

First, to improve the inhibitory activity of AcrIIA4 towards the VPR-dCas9, we increased their binding affinity by fusing to the two proteins, two orthogonal synthetic Coiled Coils (CCs) domains [29, 30] that act as “molecular magnets” promoting their dimerisation, as shown in Figure 4.a. This modification led to a 600-fold suppression of the VPR-dCas9-N8 activity, marking a threefold increase in inhibition compared to the original Anti-CRISPR/VPR-dCas9 pair without CCs, as illustrated in Figure 4.b and Supplementary Figure S3. These results hold true also in a different cellular model (Supplementary Figure S3).

**Figure 4:**
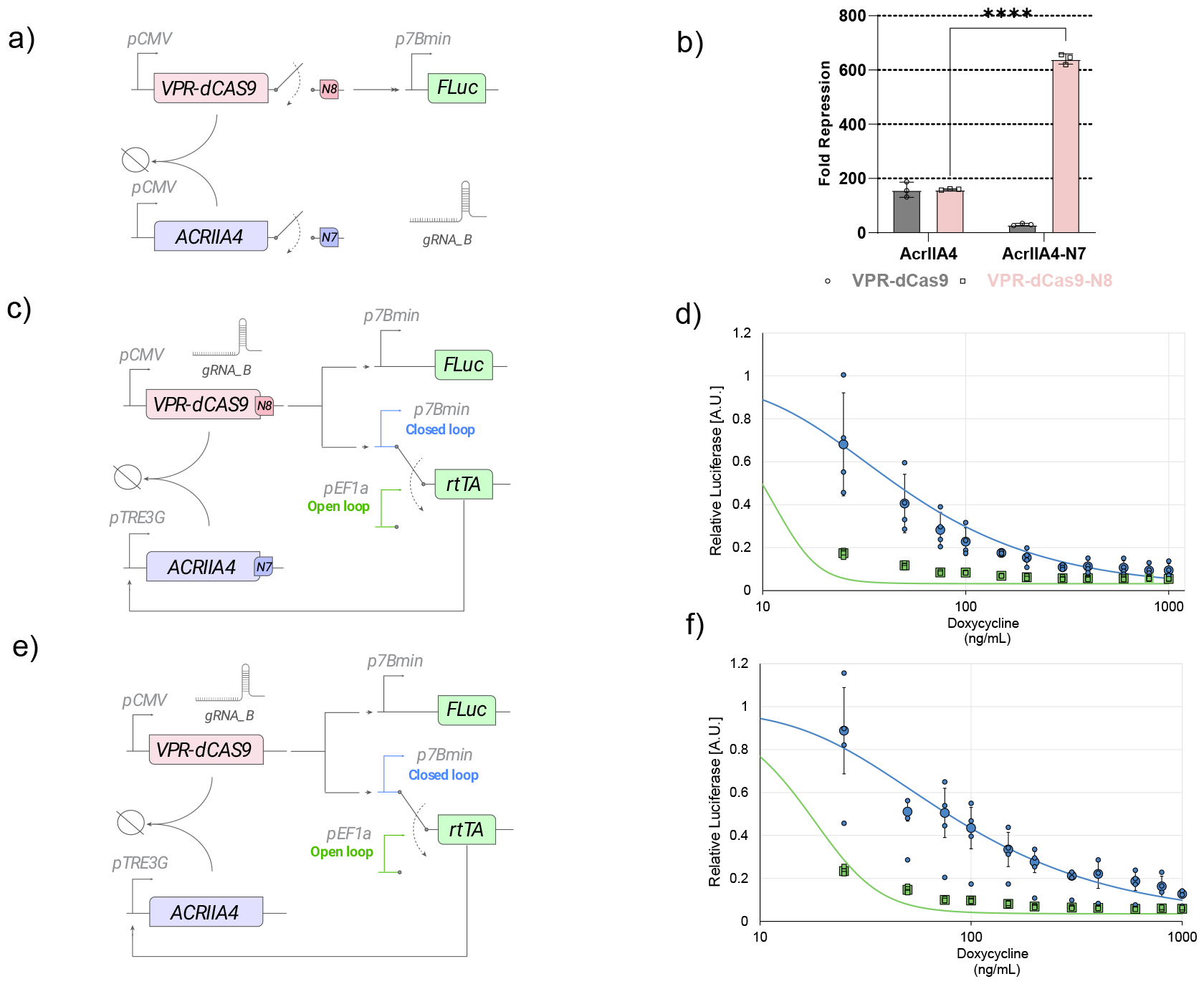
Experimental implementation of the NFL motif. **a**, Schematic representation of the constructs transfected in HEK293T cells to assess the transcriptional activity of VPR-dCas9 and AcrIIA4, with or without the cognate coiled-coiled (CC) domains N7 and N8. The promoter p7B Min in the presence of the guide RNA gRNA_B drives expression of the firefly luciferase (FLuc). **b**, Fold-repression in luminescence when co-transfecting either the VPR-dCas9 (grey bars) or the coiled-coiled fusion variant VPR-dCas9-N8 (pink bars), in presence of AcrIIA4, or of the coiled-coiled fusion variant AcrIIA4-N7. Fold repression is computed by dividing the luminescence of FLuc in the absence of the AcrIIA4 (or AcrIIA4-N7) by the value in its presence. Statistical significance is computed via a two-population t-test. **c**, Implementation of the NFL motif. The VPR-dCas9-N8 is constitutively expressed from the pCMV promoter and, in the presence of the gRNA_B, binds to the 7B pMin promoter driving the rtTA transcription factor. In the presence of doxycycline, rtTA binds to the pTRE3G promoter upstream of the AcrIIA4-N7. The VPR-dCas9-N8 is inactivated by the AcrIIA4-N7, thus closing the loop. The firefly luciferase (FLuc) under the control of the p7B_Min promoter is used as a reporter. In the open loop configuration, the rtTA is under the control of the constitutive promoter pEF1*α*. **d**, Dose-response curve to doxycycline of the NFL (blue circles) and of the open loop control (green squares). Relative Luciferase is the FLuc luminescence divided by its value in the absence of Doxycycline. The solid lines are the result of the numerical simulation of a mathematical model of the NFL. **e**, Experimental implementation of the NFL without the N7 and N8 coiled-coils. **f**, Dose-response curve to doxycycline of the NFL without CCs (blue circles) and of the open loop control (green squares). *n=4 biological replicates. A minimum of n=3 when one of the measurements was identified as an outlier (Grubbs’ test, alpha=0*.*2). Statistics analysis has been conducted through a two-way ANOVA test. **** P* ≤ *0*.*0001* ***VPR-dCas9-N8***: *nuclease-deficient Cas9 fused to the transactivation domain VPR and to synthetic coiled-coil N8;* ***AcrIIA4-N7***: *Anti-CRISPR protein fused to the synthetic coiled-coil N7;* ***rtTA***: *reverse tetracycline TransActivator 3G;* ***pTRE3G***: *Tetracycline Responsive Element promoter 3G*.

We then implemented the NFL motif as shown in Figure 4.c and detailed in Supplementary Material. Briefly, the pCMV promoter drives constitutive expression of the VPR-dCas9-N8 that acts as the *X* of the NFL design. In the presence of the constitutively expressed gRNA_B, the VPR-dCas9-N8 binds to the p7B_Min promoter upstream of the reverse-tet-TransActivator (rtTA) gene, acting as species *Z*. The rtTA protein, in the presence of Doxycycline, binds to the pTRE3G promoter and drives the expression of the modified anti-CRISPR (AcrIIA4-N7), acting as species *Y*. This protein in turn binds to and inhibits VPR-dCas9-N8, thus closing the negative feedback loop. The FLuc is also under the control of the p7B_min promoter thus tracking the expression of species *Z* and serves as the reporter. As an experimental control, we also built an “open loop” version of the circuit, as shown in Figure 4.c, where the rtTA is under the control of the constitutive promoter pEF1*α*, thus disabling the negative feedback loop.

We then measured the FLuc luminescence at increasing concentrations of Doxycycline, as reported in Figure 4.d. The dose-response curve of the NFL (blue circles) shows a gradual decrease in luminescence, in close agreement with numerical simulations of a mathematical model of the circuit (blue line). On the contrary, in the open loop configuration (green squares), there is a sharp drop in luminescence even at the lowest concentration of doxycycline, again in agreement with mathematical modelling (green line). This effect is caused by the rtTA-mediated expression of the anti-CRISPR protein in the presence of doxycycline, which binds to and inhibits the VPR-dCas9-N8, thus preventing it from activating the p7B_min promoter upstream of the luciferase. In the NFL setup, however, reduced VPR-dCas9 activity also lowers rtTA production, leading to a reduction in anti-CRISPR protein, hence resulting in a more gradual dose-response curve.

Finally, to assess the effect of the CCs domains, we built an additional version of the NFL, but this time using the original variants of VPR-dCas9 and AcrIIA4, without CCs, as shown in Figure 4.e. The dose-response curve of the NFL in Figure 4.f exhibits higher luminescence than the original NFL (with CCs) for all doxycycline concentrations (blue circles), indicative of a weaker inhibitory action of the AcrIIA4 on the VRP_dCas9, as expected. The dose response of the open loop control (green squares) is also reported in Figure 4.f.

Overall these results confirm that the circuit in Figure 4.c behaves as a NFL. Hence, we decided to implement the NFL copper biosensor by simply swapping the pCMV promoter upstream of the VPR-dCAS9-N8, for the copper-sensitive pMRE promoter, as shown in Figure 5.a. We then measured the dose-response curve for increasing concentrations of copper dichloride in the presence of a fixed saturating concentration of Doxycycline, as reported in Figure 5.d. In agreement with the quantitative modeling, the results in Figure 5.g demonstrate that the NFL motif delivers significant improvements in terms of linearity and operation range when compared to the basal biosensor. However, this comes at the cost of a worsening in leakiness and in fold change.

**Figure 5:**
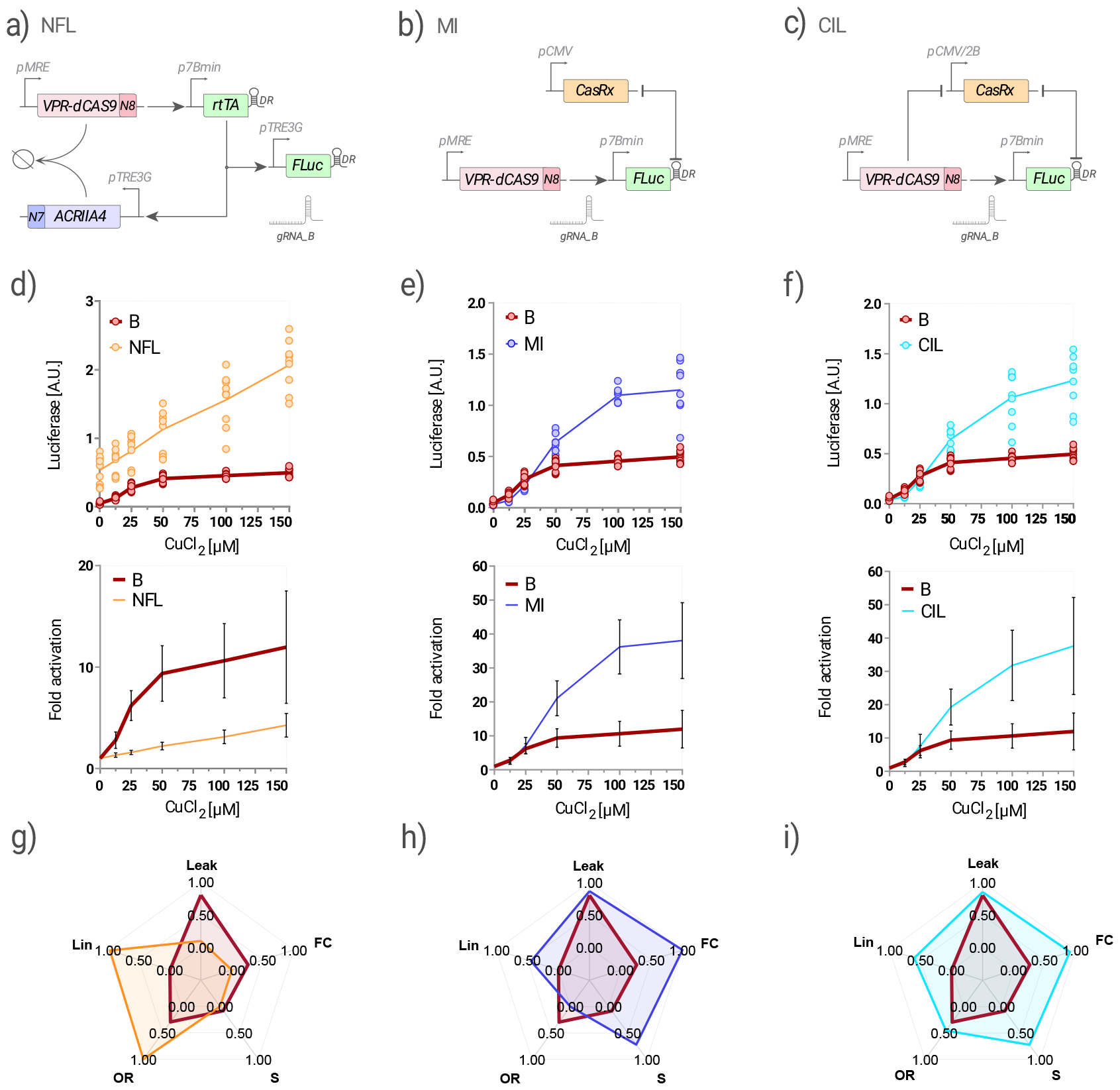
Experimental implementation of the NFL, MI and CIL biosensors. **a**, The Negative Feedback Loop (NFL) copper biosensor. **b**, The Mutual Inhibition (MI) copper biosensor. **c**, The Coherent Inhibitory Loop (CIL) copper biosensor. **d-f**, Dose-response curve of the indicated biosensor against the basic biosensor B (red line) for increasing copper dichloride concentrations. n=9 biological replicates from three independent experiments. Firefly luciferase (FLuc) expression was evaluated by luminescence measurements normalised to Renilla firefly (RLuc) luminescence and reported either as Arbitrary Units [A.U.] (upper panel), or as Fold Activation obtained by dividing each data point by the average luminescence in the absence of copper (lower panel). **g-i**, Radar plot reporting the values of the five features of each biosensor against the basal biosensor (red line). Leak: leakiness, FC: fold change, S: sensitivity to input, OR: operation range, Lin: linearity. The score ranges from 0, which is assigned to the biosensor with the worst performance, to 1 for the system with the best performance. ***CasRx***: *Cas13 RNA endonuclease RfxCas13d;* ***DR***: *direct repeat sequence cleaved by CasRx;* ***pCMV/2B***: *modified CMV promoter with two sequences complementary to gRNA_B flanking tha TATA box*.

### Experimental implementation of the MI and CIL biosensors

The experimental implementations of the MI and CIL biosensors are reported in Figure 5.b,c. They were obtained by modifying their original implementations [16]. Specifically, the VPR-dCas9-N8 protein, driven by the pMRE promoter, acts as species *X*. The CasRx endoribonuclease[31] serves as species *W*, while Firefly Luciferase (FLuc) represents species *Z*. In these two motifs, CasRx recognises and cleaves a cognate Direct Repeat (DR) sequence inserted in the 3’UTR of the FLuc mRNA. This cleavage removes the polyA signal thus leading to mRNA degradation. Since the CasRx remains bound to the DR sequence, it is unable to recognise and cleave additional FLuc mRNAs; hence, a mutual inhibition is established between the CasRx and the FLuc mRNA, as previously reported [16]. In the MI implementation, depicted in Figure 5.b, the CasRx is constitutively expressed, while in the CIL implementation in Figure 5.c, the CasRx is driven by the synthetic promoter pCMV/2B. This synthetic promoter can be repressed by VPR-dCas9-N8 in the presence of gRNA_B thanks to the presence of two DNA binding motifs downstream of the TATA box causing steric hindrance, as described in Supplementary Figure S4.

The dose-response curve for increasing copper dichloride concentrations is shown in Figure 5.e,f for both the MI and CIL motifs. Quantification of the features in Figure 5.h,i demonstrate a substantial improvement in sensitivity and fold-change, in agreement with the in silico results, a modest improvement in linearity, and no worsening in leakiness, despite the maximum expression of FLuc in the presence of copper being more than two-fold compared to the Basic copper biosensor (Figure 5.e,f). The operation range, however, is worse in the MI, as compared to the basic copper biosensor, and is unchanged in the CIL. Overall these experimental results demonstrate that the improvements provided by the MI and CIL motifs are complementary to those obtained with the NFL.

### Experimental implementation of the NFL-MI and NFL-CIL biosensors

The experimental implementation of the NFL-MI network motif is reported in Figure 6.a. The NFL motif was augmented with the constitutively expressed CasRx protein, and with the DR sequence in the 3’UTR of both the rtTA and FLuc genes. This strategy enables the CasRx to repress the expression of both genes in order to enforce minimal leakiness.

**Figure 6:**
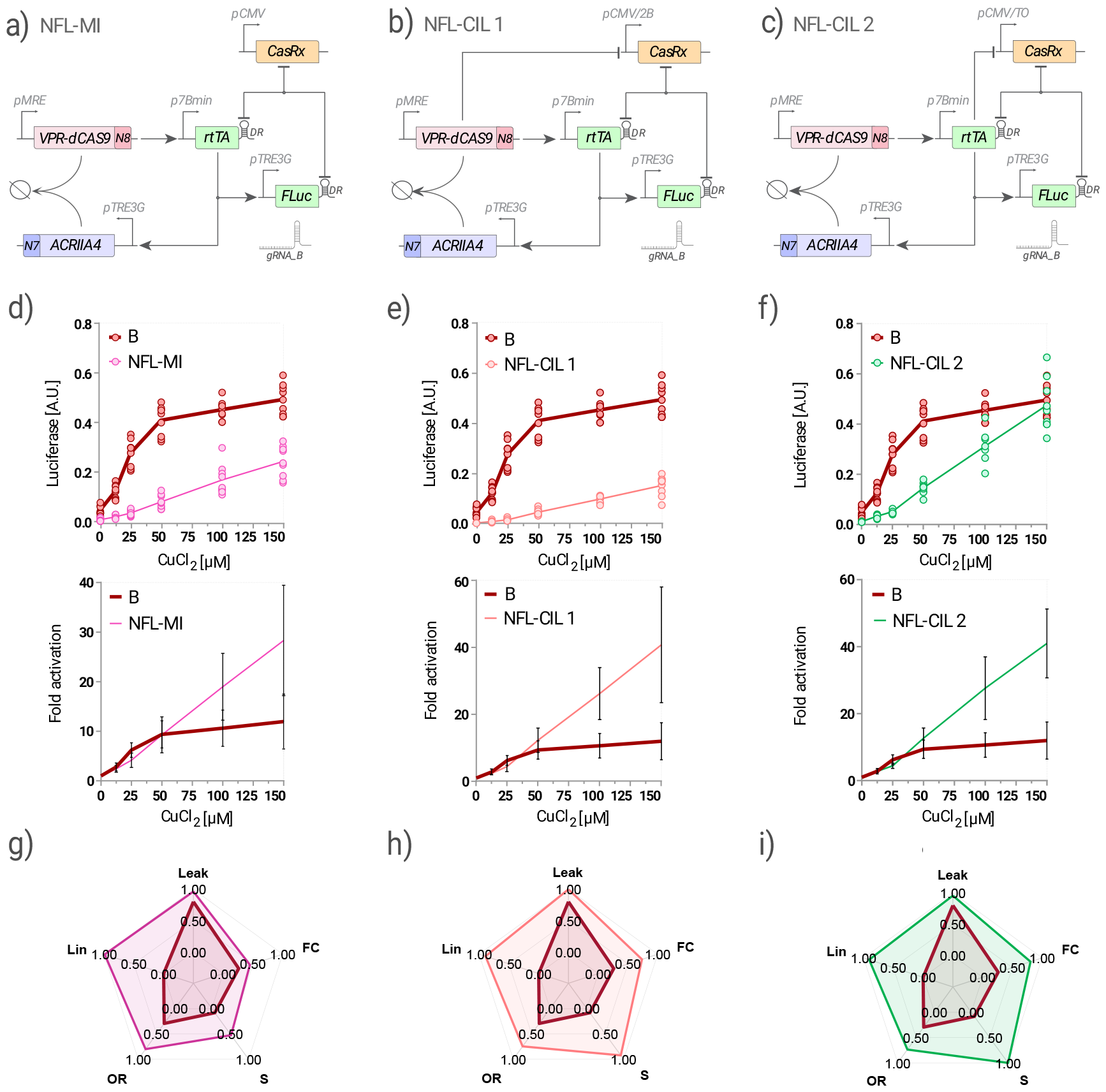
Experimental implementation of the NFL-MI, NFL-CIL 1 and NFL-CIL 2 biosensors. **a**, The NFL-MI copper biosensor. **b**, The NFL-CIL version 1 copper biosensor. **c**, The NFL-CIL version 2 copper biosensor. **d-f**, Dose-response curve of the indicated biosensor against the basic biosensor B (red line) for increasing copper dichloride concentrations. n=9 biological replicates from three independent experiments. Firefly luciferase (FLuc) expression was evaluated by luminescence measurements normalised to Renilla firefly (RLuc) luminescence and reported either as Arbitrary Units [A.U.] (upper panel), or as Fold Activation obtained by dividing each data point by the average luminescence in the absence of copper (lower panel). **g-i**, Radar plot reporting the values of the five features of each biosensor against the basal biosensor (red line). Leak: leakiness, FC: fold change, S: sensitivity to input, OR: operation range, Lin: linearity. The score ranges from 0, which is assigned to the biosensor with the worst performance, to 1 for the system with the best performance.

We then implemented the NFL-CIL version 1 network motif, as shown in Figure 6.b by swapping the constitutive pCMV promoter driving the CasRx in the NFL-MI network, for the pCMV/2B promoter that can be repressed by the VPR-dCas9-N8. Analogously, we implemented the NFL-CIL version 2 network motif as shown in Figure 6.c, by modifying the NFL-MI network, but this time swapping the constitutive pCMV promoter driving the CasRx for the pCMV/TO promoter [16, 32], which can be repressed by the rtTA trancription factor.

To derive the dose-response curves of these three biosensors, we treated Hek293T cells transfected with the relevant plasmids, at increasing concentrations of copper dichloride and measured luminescence, as done for the previous biosensors. Experimental dose-response curves are reported in Figure 6,d-f and show a substantial improvement of all the five features, as quantified in Figure 6g-i. Specifically, the NFL-CIL version 2 variant exhibited the best overall performance, again in agreement with quantitative modeling. We thus named this biosensor configuration the CASense.

## Conclusion

Here, we relied on key theoretical results from Control Engineering to design biomolecular circuits yielding high-performance biosensors in terms of leakiness, fold change, sensitivity to input, operation range, and linearity of the dose-response curve. The biosensor is abstracted as an input-output system, with the input being the specific analyte under investigation, and the output quantified through the expression level of a reporter protein.

Quantitative modeling was used to investigate the properties of eight distinct biomolecular circuits incorporating different combinations of gene network motifs endowing the system with the desired properties. Through both analytical and numerical analyses, we established that the NFL motif acts orthogonally to the MI and CIL motifs in terms of improving biosensor features. Specifically, the NFL motif confers increased linearity and operation range, whereas leakiness, sensitivity and fold-change are improved by the MI and CIL motifs.

We then experimentally implemented the biomolecular circuits by employing the flexibility of the DNA endonuclease dead-Cas9 (dCas9) as a synthetic transcription factor, and the properties of the Cas13d (CasRx) RNA endonuclease to regulate mRNA expression post-transcriptionally, to implement the gene networks and evaluated their performances against a conventional biosensor [15]. The experimental results confirmed the orthogonal properties of the gene network motifs that were inferred by the in silico analysis. Hence, by combining the NFL and CIL motifs, we succeeded in developing novel biomolecular circuits that act as high-performance biosensors. Specifically, the CASense configuration closely approximates an ideal biosensor exhibiting minimal leakiness with no reduction in the maximum reporter expression, a linear dose-response curve, high sensitivity and high fold-change. These features enable measurements of an analyte of interest, such as intracellular copper, by simply measuring the output response (i.e. luminescence).

We chose this specific example because of the critical roles that copper plays in human health and disease. Copper-related disorders include Wilson’s disease (WD) [33], a rare genetic disorder caused by defects in ATP7B copper-transporter, and Menkes disease where copper absorption is impaired by mutations in the ATP7A gene. Abnormal copper levels can also be associated with neurological disorders such as Alzheimer’s disease, Parkinson’s disease, and other neurodegenerative conditions. Measurement of copper concentration requires sophisticated analytical techniques due to the low concentrations involved, moreover there are currently no techniques able to precisely measure intracellular copper levels in living cells[34]. Therefore, developing a biomolecular copper biosensor capable of accurately measuring intracellular copper concentrations *in vitro* and *in vivo* over time could be a game changer to set-up high-throughput drug screenings and to monitor gene therapy in animal models[35, 36].

The current experimental implementation has some limitations. Specifically, the constructs required to implement the CASense are quite large thus hindering their packaging in viral vectors and limiting in vivo applications. Moreover, the CASense circuit may have unintended effects on primary cells and tissues, such as altering metabolism and stress response, as they require the integration of multiple CRISPR-CAS systems with potential off-targets.

Our newly devised biomolecular circuits offer a general strategy to design high-performance biosensors for any analyte or cellular state of interest, provided there is an existing analyte-sensitive promoter. Moreover, our study underscores the value of a theory-driven quantitative approaches in designing biomolecular circuits with desired input-output properties even in complex systems such as mammalian cells.

## MATERIALS AND METHODS

### Plasmid Construction

Most of the plasmids were constructed using the Golden-Gate based EMMA assembly kit [37], following authors’ protocol [38], NEBridge Golden Gate Assembly BsaI-HFv2 and BsmBI-v2 kits (NEB) employing custom fusion sites, while in other cases the plasmids have been constructed using the Gibson assembly method [39]. All the sequences regarding N7 and N8 coiled coils [30], used to modify the CRISPR-antiCRISPR system, AcrIIA4 [27] and FKBP-derived DD [40], gRNA inducible constructs with 7 binding sites for gRNA_B [26] and 1 or 10 binding sites for gRNA_AB [29], gRNAs sequences [29, 26], VPR-dCas9 [41], CASRx [31] and the copper – responding MRE promoter [42] have all been taken from published papers. The Tet-On® 3G system, comprising the TRE3G promoter and the rtTA protein, has been acquired from Takara Bio.

### Cell culture and transfection

The HEK293T cell line (ATCC) was cultured in DMEM Gluta-max (Gibco) supplemented with 10% Tet-Free Fetal Bovine Serum (Euroclone) and 1% Penicillin-Streptomycin (Euroclone), while the HeLa cell line (ATCC) was cultured in DMEM (Gibco) supplemented with 10% Tet-Free Fetal Bovine Serum (Euroclone), 1% L-Glutamine (Euroclone) and 1% Penicillin-Streptomycin (Euroclone). Both the cell lines have been kept at 37°C in a 5% CO2 environment. For luciferase experiment, 1,5×10^4^ (HeLa) and 2×10^4^ (HEK293T) cells per well were seeded in CoStar White 96-well plates (Corning) to perform standard transfection, while 4,5×10^4^ (HEK293T) were seeded when performing reverse transfection. For Flow Cytometry assay the same numbers of cells have been seeded in 96-well cell culture plates (Corning). After 18 hours of seedling, for standard transfection, or immediately after seedling, for reverse transfection, cells have been transfected using a home-made solution of PEI (MW 25000, Polysciences, stock concentration 0,324 mg/ml, pH 7.5) using from 200 to 250ng of DNA per well. For HEK293T cells, 5 *µ*l of PEI solution for each µg of DNA were used, while for HeLa 4 *µ*l of PEI solution for each µg of DNA were used.

### Luciferase assay

Firefly Luciferase and Renilla Luciferase expression were measured following the protocol of Dual Luciferase Assay (Promega) on a Glomax Explorer plate reader (Promega). The cells were collected 48 hours after transfection and lysed with 5X Passive Lysis Buffer (Biotin) diluted in water. Firefly Luciferase Arbitrary Units (Luciferase [A.U.]) were calculated by normalizing each sample’s Firefly Luciferase activity to the constitutive Renilla (pRL-TK, Promega) activity detected in the same sample. Relative Luciferase [A.U.] values have been calculated by dividing each sample’s Luciferase [A.U.] by the Luciferase [A.U.] of the sample with no Doxycycline induction, in the case of Doxycycline dose-response curves. The Fold Activation has been obtained by dividing the average of each data point of Luciferase [A.U.] by the average of Luciferase [A.U.] in the absence of copper. For the characterization of the NFL Motif, a Firefly Luciferase with two destabilization sequences has been used (Luc2CP, Promega), while for the biosensing experiments, a stable Firefly Luciferase (Luc2, Promega) has been used in order to compare it with the original biosensor.

### Drug treatment

Cells were treated with Doxycycline (Clontech), Shield1 (MedCehmExpress) and/or Copper dichloride (Sigma – Aldrich) immediately before transfection. Doxycycline and Copper dichloride were dissolved in H2O, while Shield1 was dissolved in DMSO. For Shield1 and Doxycycline, drugs’ concentrations are referred to the medium volume before adding the transfection mix, while for Copper dichloride, drugs’ concentrations are referred to the medium volume after adding the transfection mix.

### CF4 Analysis

Hek293T cells were plated in a 24 well and treated with increasing concentrations of CuCl2 the following day. Forty-eight hours after CuCl2 treatment, the cells were treated with 1.5 *µ*M CF4 diluted in medium, incubated for 15 minutes at 37°C, then resuspended, washed with PBS (Euroclone) and analyzed by flow cytometry[24]. Flow Cytometry analysis were carried out using an Accuri C6+ (BD Biosciences), analyzing 10000 cells for each sample. A 488-nm laser with a 670 nm LP filter was used to excite and detect CF4.

### Mathematical Modelling

We developed deterministic Ordinary Differential Equation (ODE) for each of the eight gene networks in accordance with the principles of the laws of mass action and Michaelis-Menten kinetics. Additional details regarding the construction and examination of these models can be found in Supplemeantaty Material. Numerical simulations were run using MATLAB (v. 2022b) with the ode15s function. The numerical steady-state value for each molecular species was taken at the end of a numerical simulation of 100 hours. The parameter values used to simulate the ODE models were generated with a Latin Hypercube Sampling Method to produce 10,000 parameter combinations, starting with nominal values reported in Supplementary Material. To accelerate the process of generating the parameter sampling and simulate the eight gene networks, we exploited the SimBiology libraries. The steady-state solutions of the ODE models were derived with Wolfram Mathematica using the command Solve and manually reshaped. Quantitative evaluation of leakiness, fold change, sensitivity to input, operation range, and linearity from both numerical simulations and experimental data was performed as described in Supplementary Material. These values range from 0 to 1, which denote the worst performance and the best performance, respectively.

